# Design of Novel Dehalogenases using Protein Large Language Models

**DOI:** 10.1101/2024.10.28.620469

**Authors:** Lucas Garcia, Willow Nauber, Camille Dahlgren, Ray Chung, Troy Reyes, Jael Santos, Maya Kapur, Monica Barsever, Matt Thomson, Alec Lourenço

## Abstract

Per- and polyfluoroalkyl substances (PFAS) are toxic compounds linked to cancers, infertility, and vaccine resistance at concentrations above 1 part per trillion. Owing to strong carbon-fluorine bonds, they are persistent in the environment, taking centuries to millennia to degrade. As part of the 2024 iGEM competition, several high school students local to the Pasadena area leveraged recently-developed bioinformatic tools and large language models to discover and design novel reductive dehalogenases predicted to degrade perfluorooctanoic acid (PFOA), a long chain PFAS whose manufacture is prohibited, yet still persists in the environment and drinking water. The team identified 68 enzymes with structural similarity to *Acidimicrobium sp. Strain A6* RdhA, the only specific known PFAS degrading enzyme in nature, expressed and refolded 5 of these enzymes in addition to 1 rational, and 3 large language model designs. These designs all have diverse sequences yet all are predicted to retain key substrate and cofactor binding pockets. These enzymes will be assayed on the ability to defluorinate PFOA.

## Introduction

Per- and polyfluoroalkyl substances (PFAS) are a group of man-made chemicals that are resistant to heat, water, and oil with a variety of applications, from use on non-stick pans to firefighting foam. However, exposure has been linked to increased cholesterol, hypertension, immune suppression, and kidney and testicular cancers[15]. Their involvement in a wide variety of negative health outcomes is thought to be due to their accumulation within cell membranes, which can alter membrane properties such as membrane fluidity and plasticity in addition to baseline toxicity[2, 16]. Current methods to degrade PFAS are costly, energy-intensive, and can produce shorter-chain PFAS byproducts, which may remain highly toxic[12].

A corrinoid iron-sulfer reductive dehalogenase, A6RdhA, was recently discovered to degrade PFOA and PFOS, which are both legacy species that the EPA recently mandated in 2024 to be less than 4 parts per trillion (ppt), the minimal detectable limit, in drinking water[8]. A partial sequence for A6RdhA was identified within Acidimicrobium sp. Strain A6, a soil-dwelling microbe that, when incubated with a PFOA or PFOS substrate, can partially defluorinate each respective compound[6]. While *in vitro* enzymatic activity has not been shown with isolated A6RdhA, Acidimicrobium sp. Strain A6 with RdhA knocked out abolishes PFOA degradation (Figure 1).

**Figure 1:**
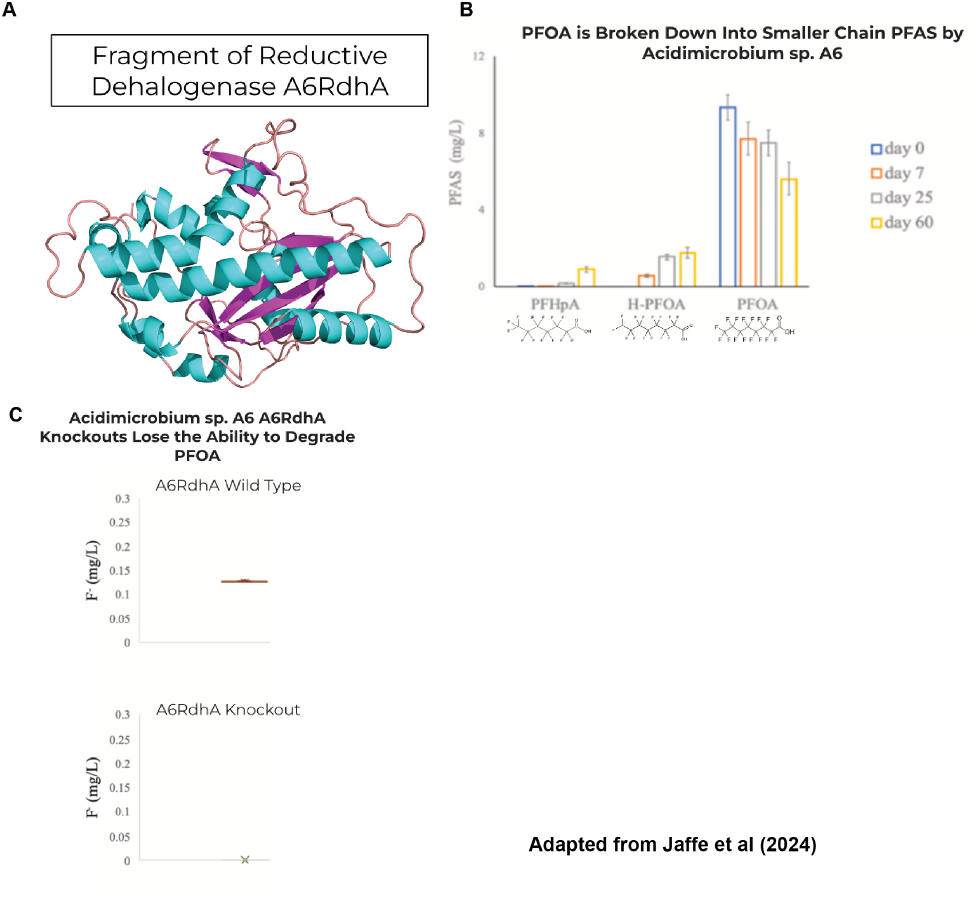
Enzyme A6RdhA is a promising starting point for enzymatic PFAS degradation. a.) Structure of enzyme A6RdhA from Acidimicrobium sp. Strain A6. b.) When Acidimicrobium sp. Strain A6 is incubated with a PFOA substrate, it is capable of partially defluorinating its structure, resulting in H-PFOA. c.) When the gene coding for the A6RdhA is knocked out, Acidimicrobium sp. Strain A6 loses its ability to degrade PFOA. These results suggest that enzyme A6RdhA is the key catalyst in this reaction.

The rarity of natural PFAS-degrading enzymes has a biophysical rationale. Fluoride is toxic to bacteria at low concentrations, with inhibitions for essential enzymes at *K*_*I*_ *<* 50*mM* in some instances [19]. Not only would a PFAS-degrading microbe need to evolve a dehalogenase, but also a fluoride ion exporter to reduce toxicity from a degradation byproduct. This dual requirement also imposes selective pressure against having a fast or efficient enzyme, meaning that there is much room for improvement through enzyme engineering.

While directed evolution has proven a successful strategy since at least the 1980s for improving protein fitness (e.g. binding strength, catalytic activity), a prerequisite to this success is the ability to screen many protein variants at once. Furthermore, natural protein sequences are often sub-optimal starting points for directed evolution, as most natural proteins are marginally stable, making the fitness landscape that protein engineers navigate through directed evolution more rugged, meaning that further mutations can more easily destabilize the proteins to the point of non-functionality[18]. Computational protein design has emerged as an alternative, but more often as a complementary approach to protein engineering. As methods for prediction of protein structure from sequence have been developed, they proved useful for protein design as well, owing to the inherent structure-function relationships embedded within. When combined with directed evolution, complete redesigns of proteins using computational methods can yield variants with multiple properties superior to the precursor. Neo2/15, an IL-2/IL-15 mimetic, was designed using Rosetta and optimized using yeast display to boast not only superior binding to the IL-2 receptor beta and gamma chain dimer, but also high expressibility in E. coli and increased thermostability upwards of 95ºC when variants were disulfide stapled[17].

Enzyme design poses additional challenges when compared with binder design, such as the requirement for precise positioning of catalytic residues and flexible backbones capable of conformational changes[11]. These features make enzyme design difficult for human designers. Large language models (LLMs), a subclass of machine learning models, have gained popularity recently for the ability to simulate human language (e.g. ChatGPT) as well as learn key features of proteins, such as protein structure[10]. Their use of unsupervised data and self-attention networks could allow these models to learn key features of enzyme design that human designers miss, provided they are trained on relevant enzyme examples from nature or otherwise.

The focus on the research was thus twofold: (1) Identify previously uncharacterized putative defluorinases from bioinformatic data and (2) use this data to train a large language model to learn principles of dehalogenase design and design novel defluorinases. We show that a partial sequence of a single PFAS degrading enzyme serves as enough of a template to generate a diverse set of structurally similar candidates. Interestingly, we find these candidates contain enough embedded information to fine-tune a large language model that learns key structural and catalytic features of reductive dehalogenases from sequence alone. We confirmed expression of several natural and generated variants as inclusion bodies and putative data suggesting they bound to the correct cofactors - key improvements toward the search and design of a defluorinase for PFAS bio-remediation.

## Results

### Identification of 68 Defluorinase Candidates

Given the existence of a single dehalogenase sequence with experimental evidence of PFAS degradation abilities, we first focused on developing a computational pipeline to identify additional candidate dehalogenases. The only sequence available for the identified PFOA degrader, A6RdhA, is from a metagenomic analysis (Genbank Accession [MK358462.1](https://www.ncbi.nlm.nih.gov/nuccore/MK358462)). Furthermore, it is a partial sequence predicted to be incapable of completing catalysis because the missing C-terminus of more than 100 amino acids makes up a large part of its active site. We posited that enzymes with similar structure to A6RdhA would have similar PFAS degrading capabilities. To find enzymes structurally similar to A6RdhA, we used the program Foldseek, which matches a query protein structure with proteins with similar structures within various protein structures and sequence databases[9].68 diverse sequences were identified after inputting the partial A6RdhA sequence as an input to the [Foldseek webserver](https://search.foldseek.com/search). The majority of these enzymes had a relatively low annotation score on [Uniprot](https://www.uniprot.org/) and were not very well characterized structurally or functionally. However, many were classified as reductive dehalogenases, which aligns with the classification of A6RdhA and fits within the family of enzymes we were researching. In addition to having high structural similarity to a known PFAS degrading enzyme, the 68 were also sequentially diverse from each other (Figure 2).

**Figure 2:**
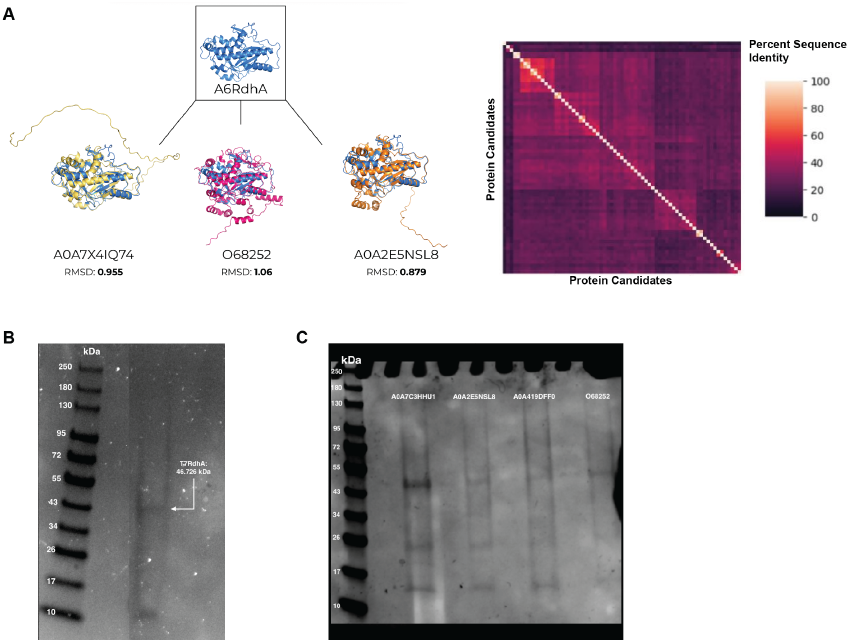
We identified, expressed, and refolded 68 putative PFAS reductive dehalogenases structurally similar to A6RdhA. a.) The partial structure of A6RdhA as a query structure to search the Foldseek database against. 68 sequences with high structural similarity, but low sequence similarity, were selected for expression and refolding. b.) T7RdhA, a previously identified corrinoid iron-sulfur protein, was expressed in *E. coli* cell free extract and refolded. c.) 4 novel variants expressed in *E. coli* cell free extract and refolded.

### Expression and Refolding of 8 Identified Enzymes

To prepare designs for experimental testing on a PFAS substrate, a subset of 8 enzymes were chosen for wetlab validation. Five of these variants were expressed in *E. coli* cell free extracts as inclusion bodies and refolded with iron, sulfur, and cyanocobalamin as previously described[13].

### Design, Expression, and Refolding of A6+T7 Chimeric Variant

Given that the only known reductive dehalogenase to degrade PFAS only has a partial sequence available, we aimed to reconstruct a functional, full length variant using homology modeling. A full-length reductive dehalogenase from bacteria *TMED77*, referred to as T7RdhA, was previously found to share sequence identity with 98% of the partial sequence obtained for A6RdhA [3]. Given the high identity between the sequences, the C-terminal region of T7RdhA was chosen to replace the missing C-terminal region of A6RdhA. The resulting variant we dubbed the A6+T7 chimera. To construct the chimera, the structures of T7RdhA and the fragment of A6RdhA were aligned, and the point at which the fragment ends was identified. The point at which T7RdhA’s sequence continues from the end of the fragment was identified. Every amino acid within T7RdhA’s sequence from where the fragment sequence ends and T7RdhA’s sequence begins was grafted onto the end of the A6RdhA fragment. This alteration of A6RdhA’s structure reconstructed its active site (Figure 3). This variant was expressed as an inclusion body from *E. coli* cell free extract and refolded with the same procedure as the natural variants (Figure 3).

**Figure 3:**
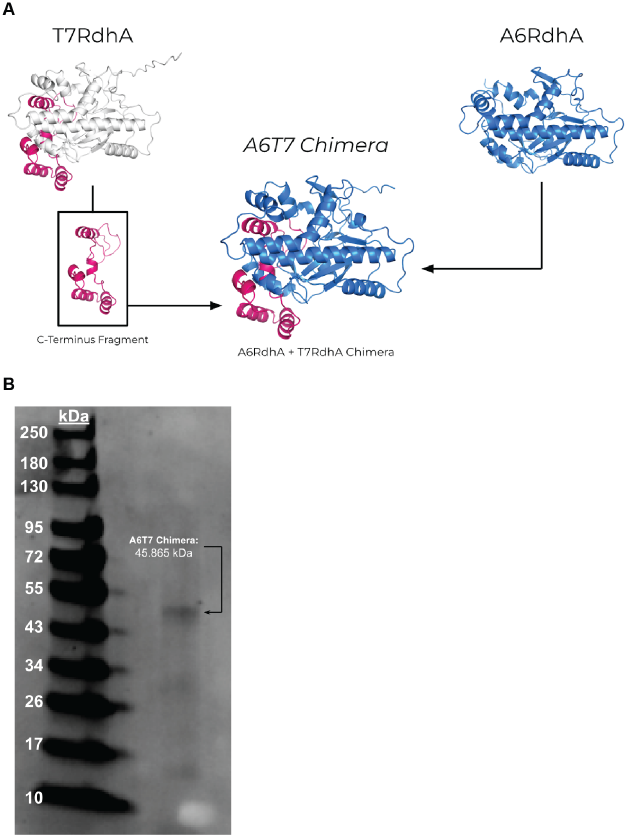
The A6+T7 Chimera reconstructed the A6RdhA fragment and was successfully expressed in a cell-free extract. a.) The A6+T7 chimera was produced by reconstructing the missing C-terminus of the original A6RdhA fragment with the C-terminus of the T7RdhA enzyme. The addition of this C-Terminus reformed the active site of the A6RdhA fragment making it more likely to be capable of binding essential cofactors. b.) The A6+T7 chimeric variant was expressed in *E. coli* cell free extract and refolded.

### Generation of Novel Dehalogenases from a Fine-Tuned Protein Language Model

To create additional variants that have the potential to degrade PFAS, we developed a generative AI model for producing reductive dehalogenases with putative defluorinase activity. To do so, we filtered the 68 enzymes identified through Foldseek to remove sequences with greater than 60 percent identity, an arbitrary cutoff to ensure adequate sequence diversity in the training data, reducing the number to 63 sequences. These were used to fine-tune a pretrained masked language model, ESM2-t33-650M-UR50D[10], to produce novel dehalogenases using an iterative unmasking process (Figure 4). These 63 sequences were converted into training data by randomly masking the sequences using a betalinear30 function[4]. As the model is further trained on selected dehalogenases, it freely generates sequences from a fully-masked prompt. After 150 epochs of training on uniquely masked repeats of the 63 enzymes, the model training is stopped and the resulting model is used for inference. The model and training files can be found [here](https://doi.org/10.5281/zenodo.13879527)[7]. 3 sequences from those generated during and after training were chosen for further analysis and expression. These sequences were found to have low sequence identity to natural sequences and to each other while retaining high structural similarity to T7RdhA (Figure 4). Additionally, when expressed in *E. coli* cell free extracts, inclusion bodies of the right molecular weight were formed and spectral scans of the reconstituted and refolded protein suggest proper binding of norpseudo-cobalamin cofactor and *Fe*_4_*S*_4_ clusters.

**Figure 4:**
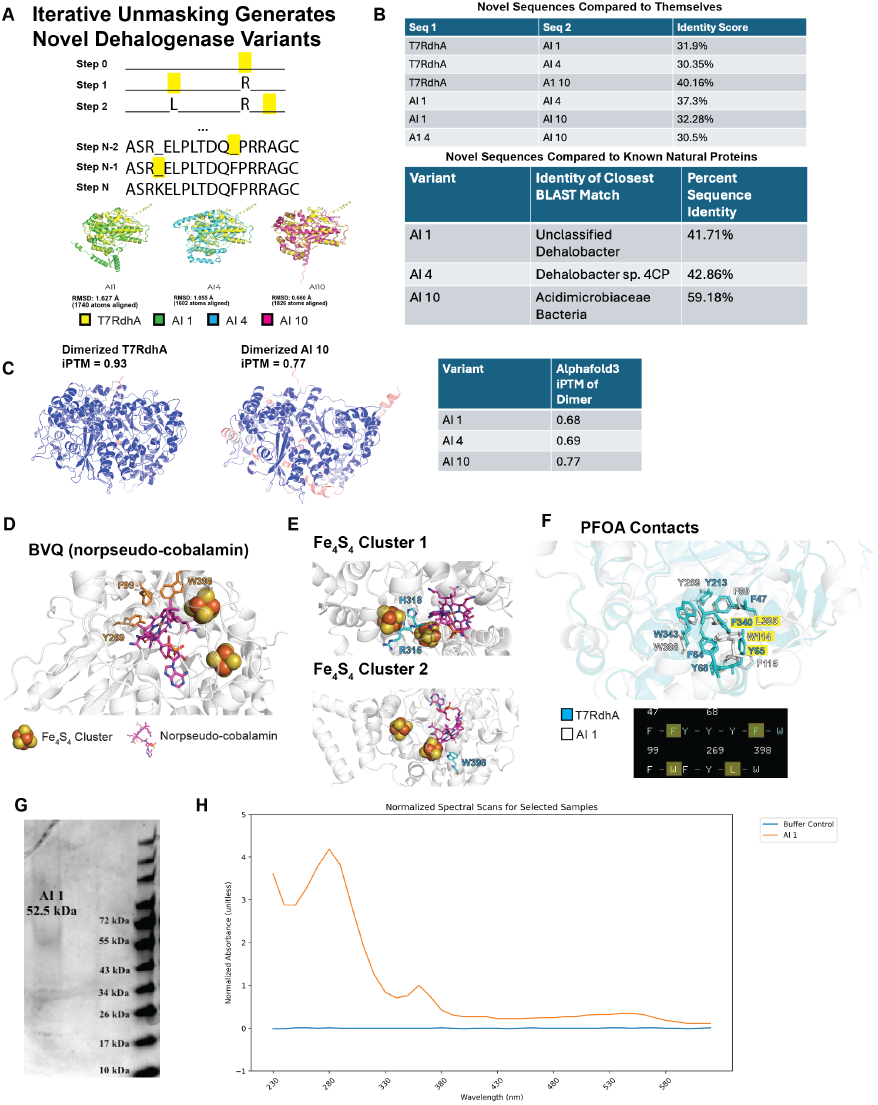
Model-generated proteins express and show evidence of co-factor binding once refolded. a.) Once trained, our masked language model generates sequences from a completely masked template sequence, randomly picking a position to unmask at each step. Three sequences from the generated set were selected based on their structural alignment to T7RdhA b.) The three variants show low sequence identity to known natural proteins and each other. c.) When cofolded with themselves in Alphafold3, the generated proteins show high interchain TM scores, predicting likely dimerization. d.) Variant AI 1 was aligned with with tetrachloroethene reductive dehalogenase containing iron-sulfur clusters and norpseudo-cobalamin (BVQ). Aligned residues in AI 1 predicted to interact with BVQ and each *Fe*_4_*S*_4_ cluster (e.) are shown. f.) Alignment of T7RdhA to AI 1 with predicted contacts to PFOA shown. AI 1 has fewer predicted contacts than T7RdhA, but many identical contacts. Different contacts (highlighted) change shape of PFOA binding pocket with minimal modification of charge. g.) We expressed AI 1 in *E. coli* cell free extract and refolded it with norpseudo-cobalamin and *Fe*_4_*S*_4_ clusters. h.) Spectral scan of AI 1 indicates norpseudo-cobalamin cofactor and *Fe*_4_*S*_4_ cluster binding, though broad peak around 550 nm indicates multiple species of reconstituted enzyme.

## Discussion

### An Integrated Approach to Protein Design

In this work, we have successfully outlined an enzyme design strategy that leverages existing tools to collect relevant training data for fine-tuning pre-trained protein large language models. By integrating existing approaches, we have circumvented the “low N” problem of having limited positive examples of enzymes in nature that perform the desired catalysis. This approach is particularly relevant to emerging focus areas such as PFAS or microplastics remediation, where there is a scarcity of confirmed natural enzymes capable of performing the desired catalytic reactions.

Our current generation process is “physics and chemistry free,” as the large language model is able to implicitly learn from a relatively small number (n=63) of natural sequences alone key structural and catalytic features of dehalogenases. We attempted to integrate an orthogonal computational screening using a combination of docking and molecular dynamics simulation, but preliminary results failed to identify key differences between candidates and clear negative control sequences. This may be exacerbated by the fact that many long-chain PFAS are known to bind to many proteins, including several in the human body, in part contributing to their bio-accumulation[5].

### Toward a PFAS Degrader

Given their status as “forever chemicals,” adverse effects from PFAS contamination can only be expected to increase in the absence of a human-made solution. Biological solutions have been hampered by the paucity of natural candidates capable of specifically degrading their carbon-fluorine bonds. Here, we have greatly expanded that set of candidates, setting up a generation and expression pipeline that can be expanded upon to further enable the search for a viable defluorinase.

## Methods

### DNA Design, Construction, and Preparation

DNA was ordered from IDT or Twist Biosciences as gBlocks or gene fragments, respectively driven by a *σ*_70_ promoter and gene 10 leader sequence containing a strong ribosome binding site[14]. A T7 terminator sequence as added downstream of each coding region. A hexahistidine tag was added to the N-terminus of all coding regions. Each dehalogenase sequence has been entered into the iGEM parts registry as IDs K5035003-K5035010. DNA was amplified with PrimeStar GxL Premix (Takara, R051) for 30 cycles using a 10 second denaturation at 98ºC, 15 second annealing at 50.9ºC, and 60 second extension at 68ºC. DNA was PCR purified using CleanNGS beads (Bulldog Bio, CNGS050).

### Cell-free Enzyme Expression and Purification

45 *µ*L of myTXTL (Arbor Biosciences, 540300) was mixed with 15 *µ*L of DNA in a 1.5 mL microcentrifuge tube and incubated at 29ºC for at least 6 hours. Following expression, a denaturing purification using 6M guanidium hydrochloride was done using Ni-NTA spin columns (NEB, S1427). Column elution was concentrated using an Amicon 30 kDa filter (Millipore Sigma, UFC503008) to concentrate the enzymes.

### Enzyme Refolding

First, 9 *µ*L of denatured enzyme was resuspended in 91 *µ*L of denaturation buffer containing 8 M urea. Next, the denatured enzyme (100 *µ*L) was mixed with 2.5 *µ*L of reduction buffer (100 mM Tris-HCl pH 7.5, 200 mM DTT) under a layer of 150 *µ*L of mineral oil, achieving a final DTT concentration of 5 mM. The mixture was incubated at room temperature for 30 minutes while stirring at 1400 rpm on a thermomixer. Following the reduction step, Fe buffer (100 mM Tris-HCl pH 7.5, 100 mM *FeCl*_3_) and S buffer (100 mM Tris-HCl pH 7.5, 30 mM *Na*_2_*S*) were added to the protein mixture, ensuring a 50-fold molar excess of Fe and S relative to the enzyme. The solution was incubated for 90 minutes, again stirring at 1400 rpm.

### UV-Visible Spectroscopy

Cofactor binding was assessed by adding 1 *µ*L of protein solution to the pedestal of a NanoDrop™ One. Absorbance was measured for wavelengths from 230-620 nm in 10 nm increments. The data was normalized by the absorbance at 360 nm.

### Alphafold Structure Predictions

All structures used for analyses were obtained via prediction with the Alphafold3 web-server[1].

### Protein large language model fine-tuning

The 68 enzymes from Foldseek were filtered by removing 5 enzymes with more than 60% identity. The resulting sequences were duplicated 150 times and masked according to the beta-linear30 function[4].The base model, *esm2 t33 650M UR50D*, was fine-tuned for 1 epoch and generated dehalogenases after every 250 training examples seen using the iterative unmasking procedure. For iterative unmasking, a sequence length from the training set was chosen at random. An amino acid is unmasked at a random, masked position, and the process is repeated until the entire sequence is unmasked. The final trained model was also used to generate candidate sequences. Sequences were chosen based on the structural similarity to T7RdhA for further computational and experimental analysis. We have included a google colab with the training code available at this [link](https://colab.research.google.com/drive/1pdSqT2oSiedgZi1dtnViZaSTR_l8cNBO?usp=sharing).

Note that this example notebook trains *facebook/esm2 t6 8M UR50D* instead of *esm2 t33 650M UR50D* to make training possible using the Google-provided free T4 GPU. Helper files and the trained model can be found [here](https://zenodo.org/records/13879527).

## Acknowledgements

We thank our extensive support community for their help in completing this work, all of whom can be listed on the IEA iGEM [team wiki](https://2024.igem.wiki/iea/gratitude) and in our [Human Practices page](https://2024.igem.wiki/iea/human-practices). Specifically, we thank Nicole Endacott and Institute of Educational Advancement for facilitating the logistics of program, including travel to the iGEM Jamboree. We thank our numerous donors, especially Linda McManus and Dr. Bill Bridges, PhD, for making travel to the Jamboree possible. We thank Dr. Mamadou Diallo, PhD, for pointing us to relevant resources related to biological PFAS degradation as well as critical discussions and feedback. We thank Melody Wu and Jason Redick for their assistance advising on various coding and iGEM specific questions. We also sincerely thank Professor Ariel Furst and Melanie Gut for their assistance with enzymatic assays as well as troubleshooting folding of iron-sulfur cluster containing proteins.

## Notes

https://2024.igem.wiki/iea/index.html

### Competing Interest Statement

The authors have declared no competing interest.

